# Divergent evolution of sleep functions

**DOI:** 10.1101/2023.05.27.541573

**Authors:** Michaela Joyce, Federica A. Falconio, Laurence Blackhurst, Lucia Prieto-Godino, Alice S. French, Giorgio F. Gilestro

**Affiliations:** Department of Life Sciences, Imperial College London, London, UK; The Francis Crick Research Institute, London, UK

## Abstract

Most living organisms have evolved to synchronize their biological activities with the earth’s rotation, a daily regulation of biology and behaviour controlled by an evolutionary conserved molecular machinery known as the circadian clock. For most animals, circadian mechanisms are meant to maximize their exposure to positive activities (*e.g.:* social interactions, mating, feeding – generally during the day) and minimize their exposure to peril (*e.g.:* predation, weather, darkness – generally during the night^1^). On top of circadian regulation, some behaviours also feature a second layer of homeostatic control acting as a fail-safe to ensure important activities are not ignored. Sleep is one of these behaviours: largely controlled by the circadian clock for its baseline appearance, it is at the same time modulated by a – poorly understood – homeostatic regulator ensuring animals obey their species-specific amount of daily sleep^2^. An evolutionary conserved homeostatic control is often considered the main evidence for a core biological function of sleep beyond the trivial one (that is: keeping us out of trouble by limiting our energy expenditure and exposure to danger^3,4^) and it is hypothesized that sleep evolved around this mysterious basic biological function. Here we characterize sleep regulation in a group of seven species of the *Drosophila* genus at key evolutionary distances and representing a variety of ecological niche adaptations. We show that the spontaneous circadian-driven aspects of sleep are conserved among all species but the homeostatic regulation, unexpectedly, is not. We uncover differences in the behavioural, cell-biological and neuro-pharmacological aspects of sleep and suggest that, in Drosophilids, sleep primarily evolved to satisfy a circadian role, keeping animals immobile during dangerous hours of the day. The homeostatic functions of sleep evolved independently, in a species-specific fashion, and are not conserved.

## Introduction and results

Studying the evolution of sleep is paramount to understand its functions, and in the past century extensive work has gone to characterize sleep in the most disparate species^5,6^, often unveiling puzzling findings. We still do not understand, for instance, how some animals – such as elephants ^7^, giraffes^8^ or cavefish^9^ – spontaneously sleep less than one or two hours a day, while others – like bats^10^ or koalas^11^ – almost completely fill their days with sleep; yet others, such as migratory birds, can adapt their sleep amount to their changing ecological needs, compressing it from many hours to mere minutes a day when the migratory instinct commands^12^. Most of the observations on the evolutionary variety of sleep and its plastic adaptation are important and insightful^13^, but conclusions are limited by the large evolutionary distance of the studied species.

Here we introduce the *Drosophila* genus as an ideal evolutionary playground to study the behavioural and genetic bases of sleep traits through a genetically well characterized group of animals at divergent evolutionary distances spanning 50+ million years of evolution. In particular, we focused our study on seven species with divergent ecological niches^14^ (Fig. 1a) and geographical ancestral origins^15–18^ (Fig. 1b). Using an automatic video tracking system^19^, we analysed spontaneous sleep in multiple independently-caught natural strains and found a qualitatively similar pattern of sleep and activity in all the species analysed (Fig. 1c and Fig. 1 supplement 1a,b). In oscillating light-dark conditions, all Drosophilids showed recognizable crepuscular peaks of activity (Fig. 1 supplement 1b,c), with their sleep mostly concentrated during the night and with male animals showing a prominent post-meridian sleep episode (Fig. 1c,e), previously identified in *D. melanogaster* as “*siesta”*^20^ and tightly bound to the circadian clock^21^. In terms of circadian regulation of sleep and activity, *D. virilis* was the only outlier in our group, showing no sexual dimorphism in the *siesta* (Fig. 1c,e) and – as previously reported^22^ – strong arrhythmicity in the absence of circadian entrainment by light (Fig. 1d). *D. virilis* is currently considered a cosmopolitan species, but its widespread localization is believed to be a very recent development in evolutionary terms, most likely linked to human movements. *D. virilis* may have originated from and adapted to the arid regions of Iran or Afghanistan from the early Miocene^23^ and this evolutionary selection may explain the unusual display of afternoon *siesta* in females (Fig. 1c-e), likely evolved as a sheltering mechanism to escape the hostile conditions of a desertic afternoon. A similar phenotype was recently observed in the closely related *D. mojavensis,* and *D. arizonae* for which it was also proposed to be a mechanism of evolutionary adaption to stressful desertic conditions^24^.

**Fig. 1.**
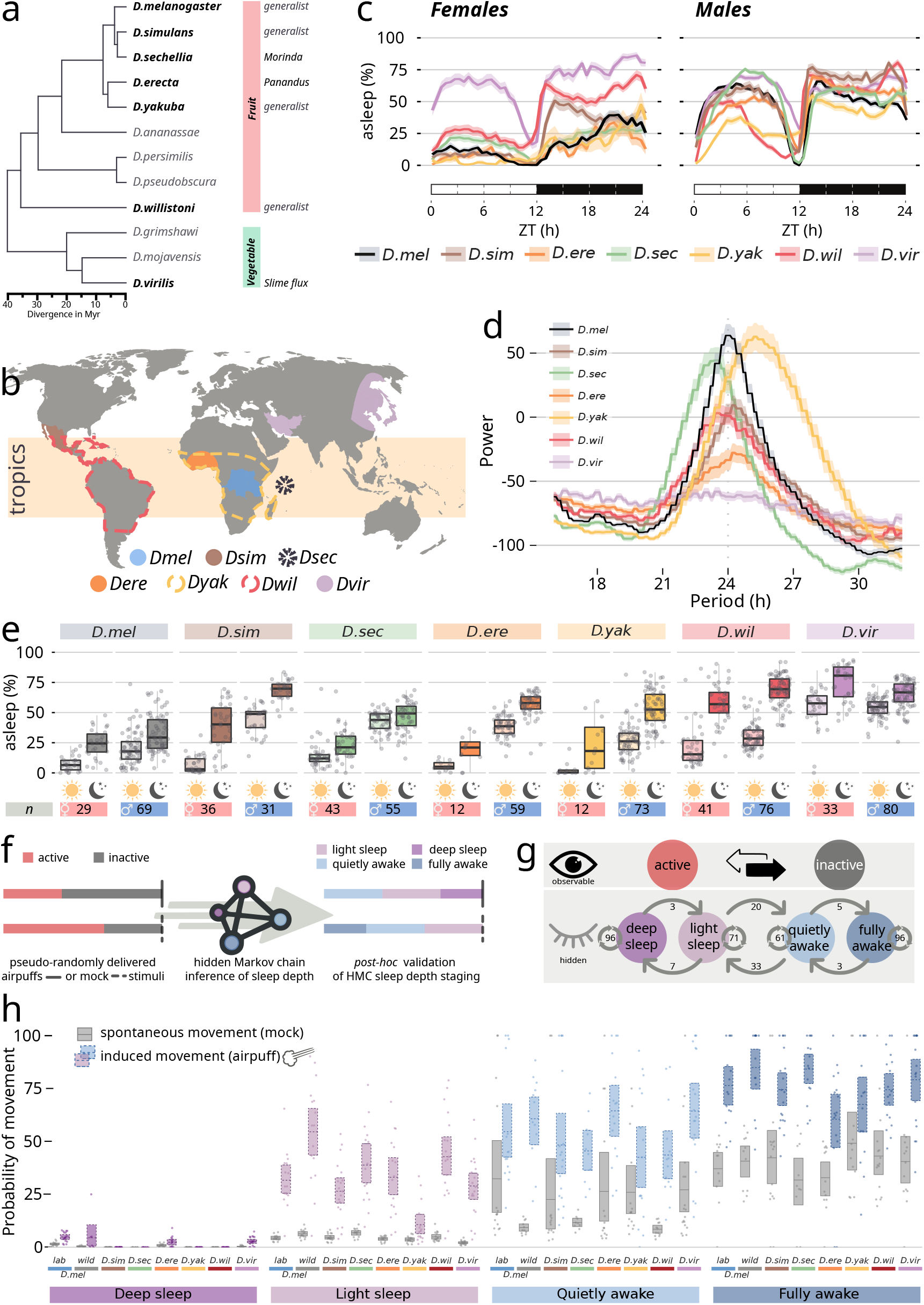
Spontaneous sleep in seven *Drosophila* species. **a,** Partial evolutionary tree of the *Drosophila* genus, highlighting in bold the seven species used in this study and their ecological niches (right). **b,** World map indicating the geographical ancestral range of each of the species employed in this study. **c,** Spontaneous, baseline sleep profile of the seven species across the 24 h period in females (left) and males (right). The white-dark bars under this all figures indicate light-dark periods. ZT: zeitgeber or relative time. **d,** power analysis of the circadian period of activity in constant dark conditions in all seven species. **e,** quantification of sleep amount during the day (ZT 0-12, sun icon) and during the night (ZT 12-24, moon icon) in females (left, pink) and males (right, blue) for all seven species. **f,** Schematics of the experimental analysis in g,h. Ethoscopes collect activity data in real time and deliver air puffs or mock stimuli after random periods of inactivity. A hidden Markov chain model trained on species-specific data classifies sleep stages. **g,** the transition parameters of the *D. melanogaster* trained model. **h,** the probability of movement following the airpuffs (coloured boxes) or the mock stimuli (grey boxes) in all seven species, by sleep stage.

**Figure 1 supplement 1.**
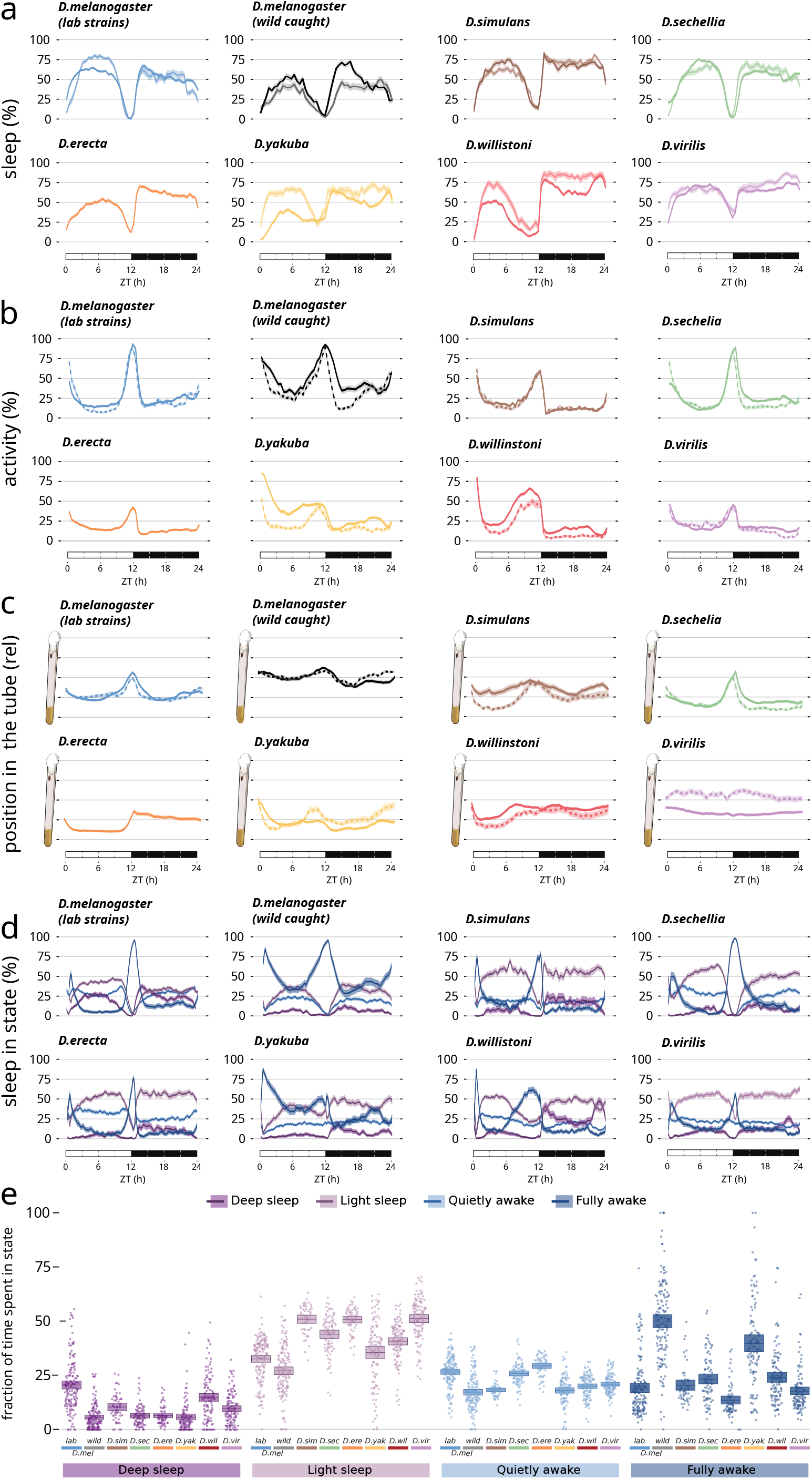
Characteristics of baseline sleep in the tested species. **a,** Quantification of sleep amount over 24 h in all tested species. With the exception of *D. erecta,* two different geographically independent strains were analysed for all species, indicated by different shades of colour. The two *D. melanogaster* laboratory strains analysed are CantonS and OregonR. **b,** Activity profile and **c,** average position inside the experimental tube over 24 h for the strains shown in a. **d,** Sleep stages in all seven species during the 24 h period. **e,** Fraction of time spent in each given state over the 24 h period in all tested species.

To investigate how profound is the evolutionary conservation of sleep in Drosophilids, we created a novel paradigm for the analysis of sleep depth. The system builds on a previously described hidden Markov chain model^25^, here complemented and validated by a robotic system^26^ able to measure arousal threshold by challenging flies with sensory stimuli delivered after random periods of inactivity (ranging from 1 to 60 minutes). Each challenging stimulus, and its consequent positive or negative response, are then collated, analysed, and correlated *ex post*^27^ (Fig. 1f-h and Fig. 1 supplement 1d,e). The combined analysis of sleep staging and arousal threshold proved valuable in three ways: i) it confirmed that the ethoscope based video tracking system could indeed reliably detect sleep in all species, even though it was initially developed for *D. melanogaster^19^;* ii) it liberated our analysis from the arbitrary definition of sleep as five minutes of immobility (also known as the five minutes rule^25,28,29^) by adopting an agnostic criterion instead; iii) it allowed us to quantify and describe the actual composition of different sleep stages during the 24 h, as well as their differences in evolution across species (Fig. 1 supplement 1d). Interestingly, we found that in all species, deep sleep mostly concentrates in the early hours of the night (ZT 12-15 – Fig. 1 supplement 1d) with *D. virilis* and *D. willinstoni* showing a second peak of deep sleep towards the end of the night. Notably, a lab cultivated strain of *D. melanogaster* (CantonS) showed an overall greater amount of deep sleep throughout the 24 h and, quite uniquely, a conspicuous amount of deep sleep during *siesta* too (Fig. 1 supplement 1d). We speculate this could be the consequence of an artificial adaptation to maintaining undisturbed sleep even in the crowded environment of a laboratory test tube.

It appears, therefore, that those aspects of sleep and activity that are meant to be regulated by the circadian clock are generally well conserved among Drosophilids but what about the second layer of regulation, homeostasis? In any animal, the easiest way to measure sleep homeostasis is to forcefully reduce sleep amount through techniques of mechanical interference to subsequently induce, and then quantify, a period of recovery sleep known as “rebound sleep”. To test rebound sleep across species we combined ethoscopes with a robotic machine able to selectively disturb the sleep of single flies by rotating their housing tube after every 30 seconds of immobility^19,30^ (Fig. 2a). To limit the amount of stress and confounding factors, the machine can recognize sleep using video tracking^19^ and will interfere exclusively with flies that are immobile for a specified amount of time, leaving the awake experimental companions undisturbed^19^. We found all seven tested species to be sensitive to this kind of mechanical interference, although with different degrees of arousal threshold (Fig. 2 supplement 1a).

**Fig. 2.**
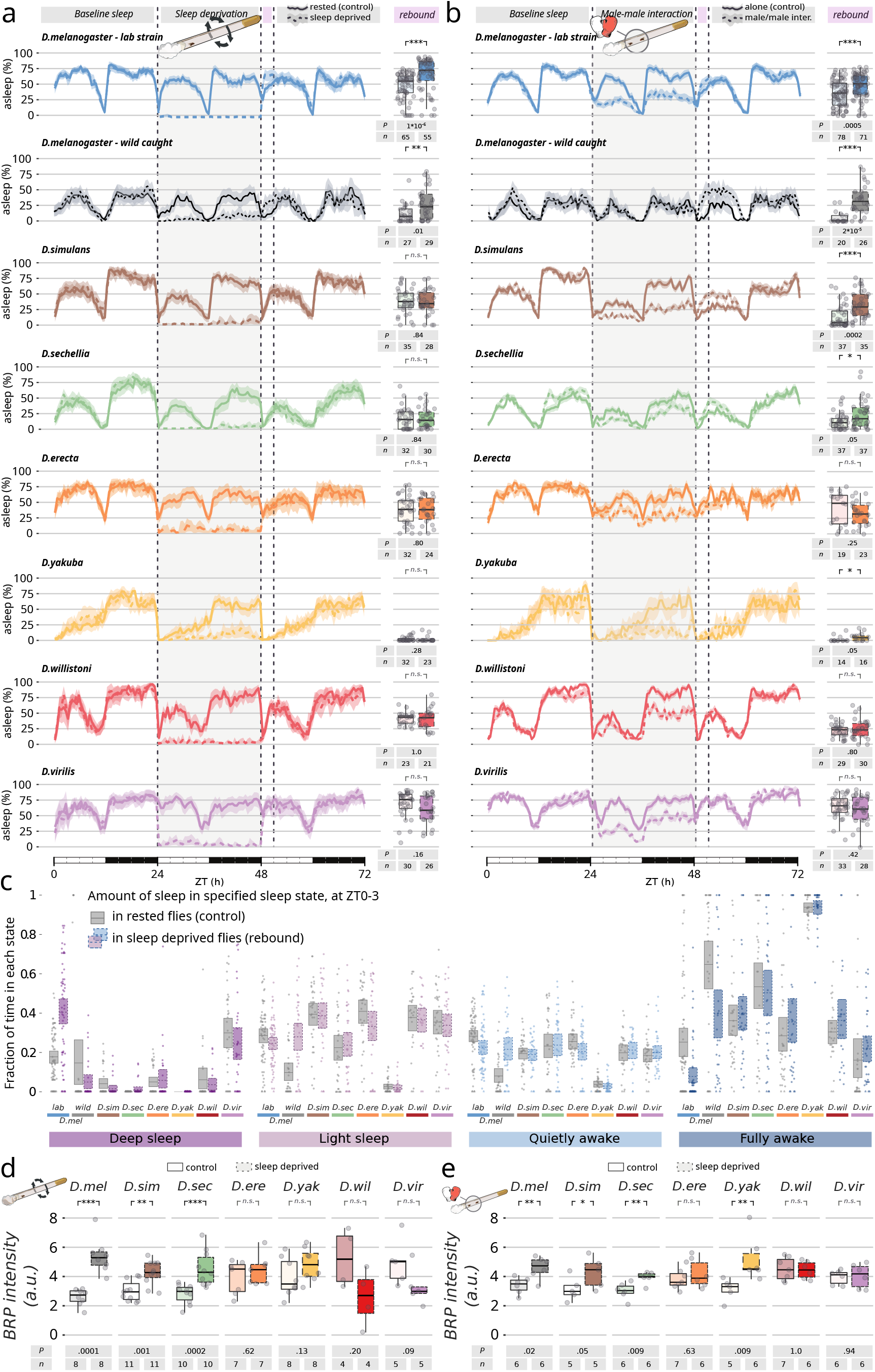
Different homeostatic rebound after mechanical or social sleep deprivation. **a-b**, 72 h sleep profile (left) and quantification of rebound (right) in flies subjected to 24 h of mechanically-induced sleep deprivation, a, or 24 h of socially-induced sleep deprivation, b. Each panel features one strain from each of the seven wild-caught species, or CantonS. In all panels, the sleep profile of rested flies is shown as a continuous line, while sleep-deprived animals are shown in a dashed line. The sleep rebound at ZT 0-3 is quantified on the right side of each sleep profile. Numbers of animals (Ns) and P-values of sleep-deprived *vs* control are shown below each panel. **c,** sleep stage analysis of all species during rebound time (ZT 0-3) after mechanical sleep deprivation. Increase in sleep depth is observed only in the laboratory strain of *D. melanogaster*. **d-e,** Quantification of the change in BRP expression in the brain of sleep-deprived flies after mechanical (**d**) or social male-male (**e**) sleep deprivation. Numbers of animals (Ns) and P-values of sleep-deprived *vs* control are shown below each panel in d and e. *** P<0.001; ** P<0.01; * P<0.05.

**Figure 2 supplement 1.**
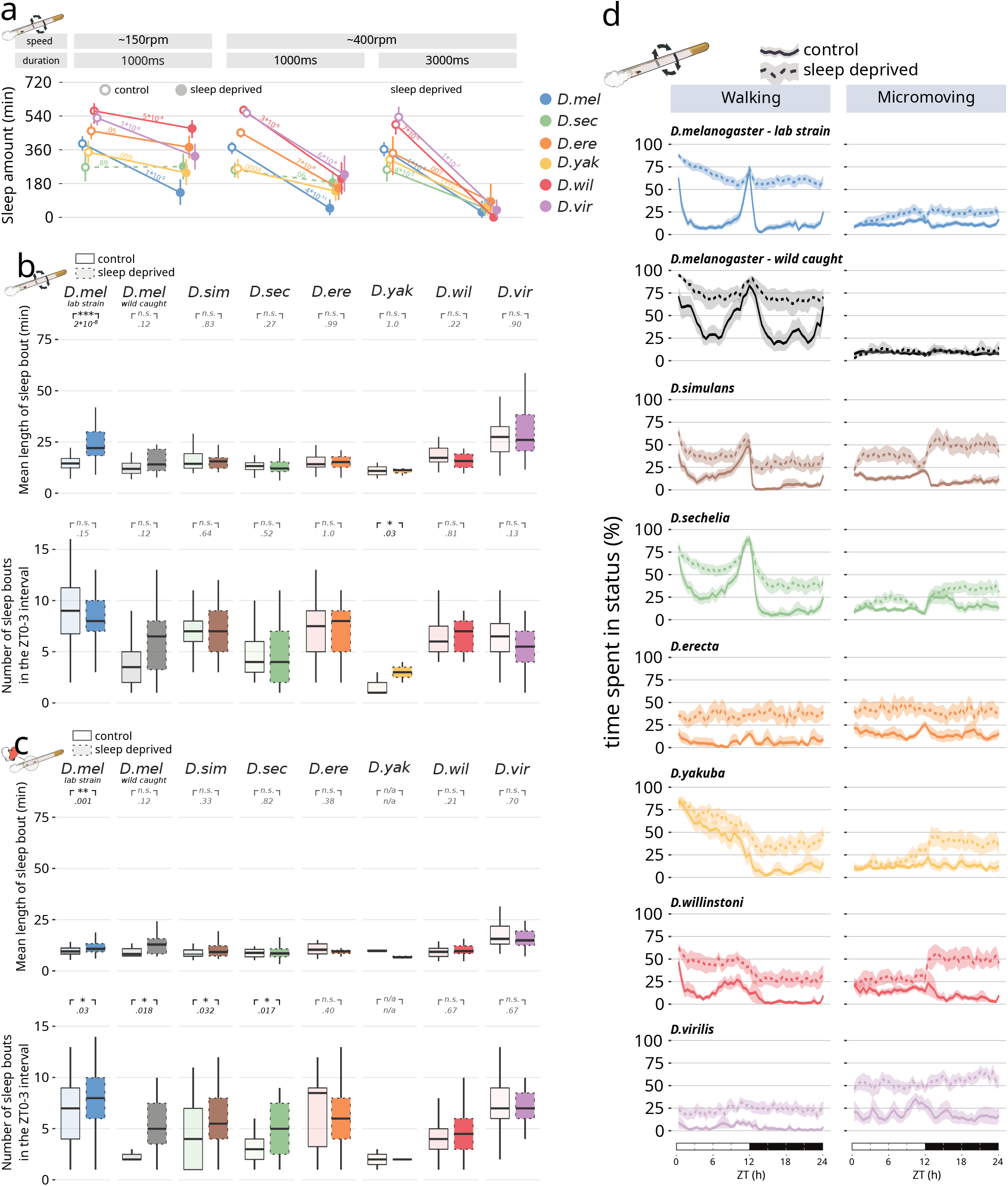
Characteristics of sleep deprivation and rebound in the tested species. **a,** Quantification of sleep in six *Drosophila* species during the sleep deprivation night using three different settings of mechanical stimuli varying speed (∼150-400 revolutions per minute) and duration (1-3 seconds). P-values are shown for each comparison. Dashed lines indicate P-values above 0.05. **b,** Mean length of sleep bouts and mean number of sleep bouts at ZT 0-3 in control (continuous lines) or animals mechanically sleep-deprived (dashed). **c,** same as b but with male-male sleep deprivation. **d,** Quantification of behaviour across the 24 h in mechanically sleep-deprived animals (dashed lines) or rested control animals (continuous line).

We then subjected all species to the most efficient stimulus (rotations of 400 rpm, 3 seconds) for 24 hours (Fig. 2a) thus forcing all the treated animals to lose most if not all of their sleep for the duration of the experiments. Surprisingly, only *D. melanogaster* showed signs of sleep rebound after mechanical sleep deprivation and no sleep rebound was detected in any of the other species, neither in terms of sheer sleep amount (Fig. 2a), nor in terms of sleep depth (Fig. 2c and Fig. 2 supplement 1b,c).

After sleep deprivation, *D. melanogaster* slept longer (and lab raised CantonS *D. melanogaster* slept deeper too – Fig. 2c) but all other species showed no signs of homeostatic rebound. Even more impressively, this difference between species persisted when sleep deprivation was pushed further to cover an uninterrupted period of 168 hours: after seven full days of continuous mechanical sleep deprivation, *D. melanogaster* showed a homeostatic rebound as previously reported^30^, but none of the other six Drosophilae did (Fig. 2 supplement 2). The technical nature of this experiment also allowed us to infer and compare the build-up in sleep pressure, which, in conditions of chronic deprivation, is a manifestation of homeostasis (Fig. 2 supplement 2). Given that our robotic ethoscopes are programmed to deliver mechanical stimuli (tube rotations) only to animals showing signs of sleep, the number of delivered stimuli over time can be used as a proxy of sleep pressure: the stronger the desire to rest, the higher the number of stimuli received^30^.

**Figure 2 supplement 2.**
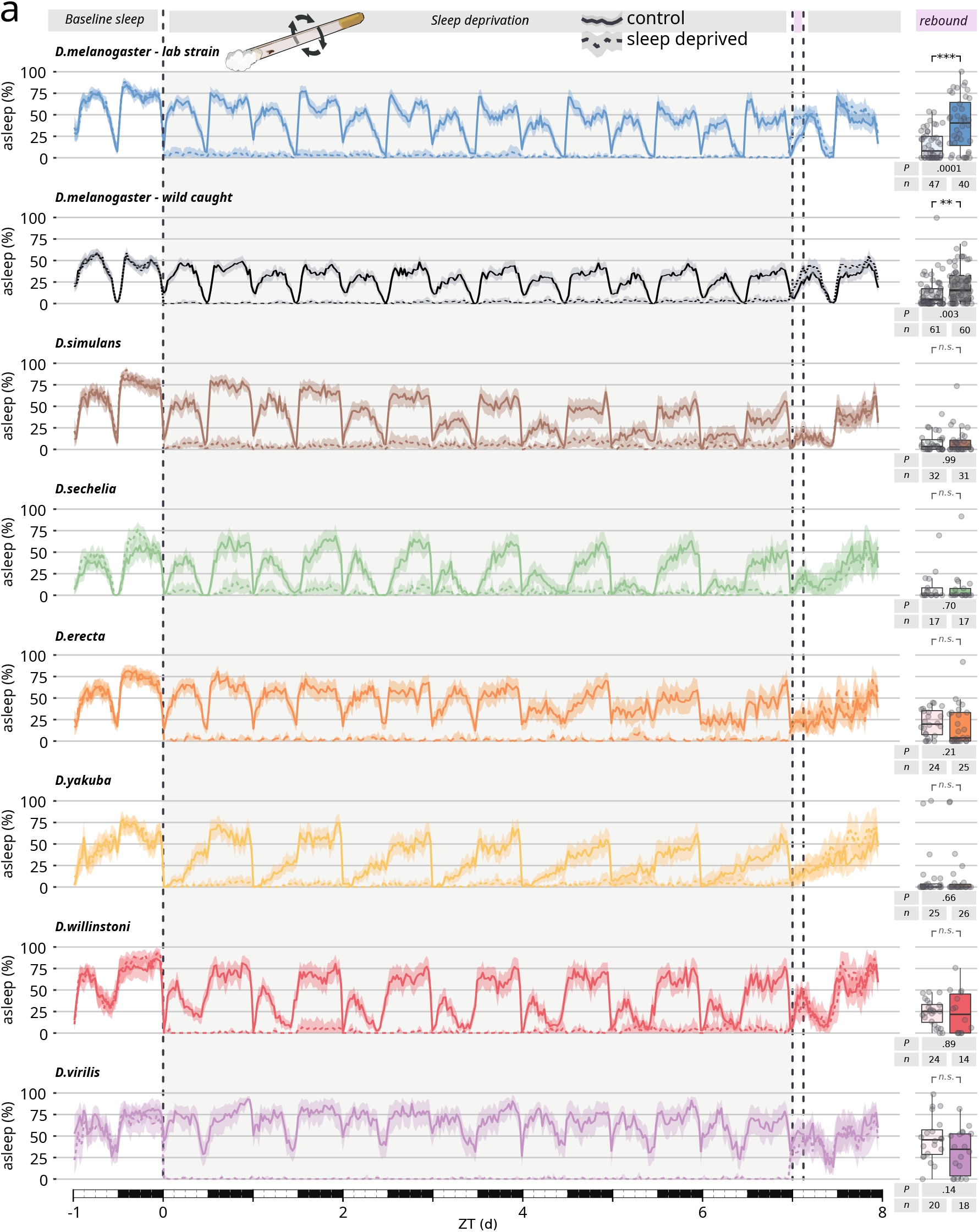
Chronic, seven days long sleep deprivation in the tested species. **a,** Total sleep profile (left) and quantification of rebound (right) in flies subjected to seven continuous days of mechanically-induced sleep deprivation. Each panel features one strain from each of the seven wild-caught species, or CantonS. In all panels, the sleep profile of rested flies is shown as a continuous line, while sleep-deprived animals are shown in a dashed line. The sleep rebound at ZT 0-3 is quantified on the right side of each sleep profile. Numbers of animals (Ns) and P-values of sleep-deprived *vs* control are shown below each panel. *** P<0.001; ** P<0.01; * P<0.05.

In line with what observed thus far, a strong week-long buildup in sleep pressure was observed only in *D. melanogaster,* whilst all other species showed no increase of sleep pressure during the seven days of sleep deprivation. If anything, almost all of them showed a decrease in sleep pressure: a possible sign of plasticity and adaptation to the sleep-deprivation treatment (Fig. 2 supplement 3).

**Figure 2 supplement 3.**
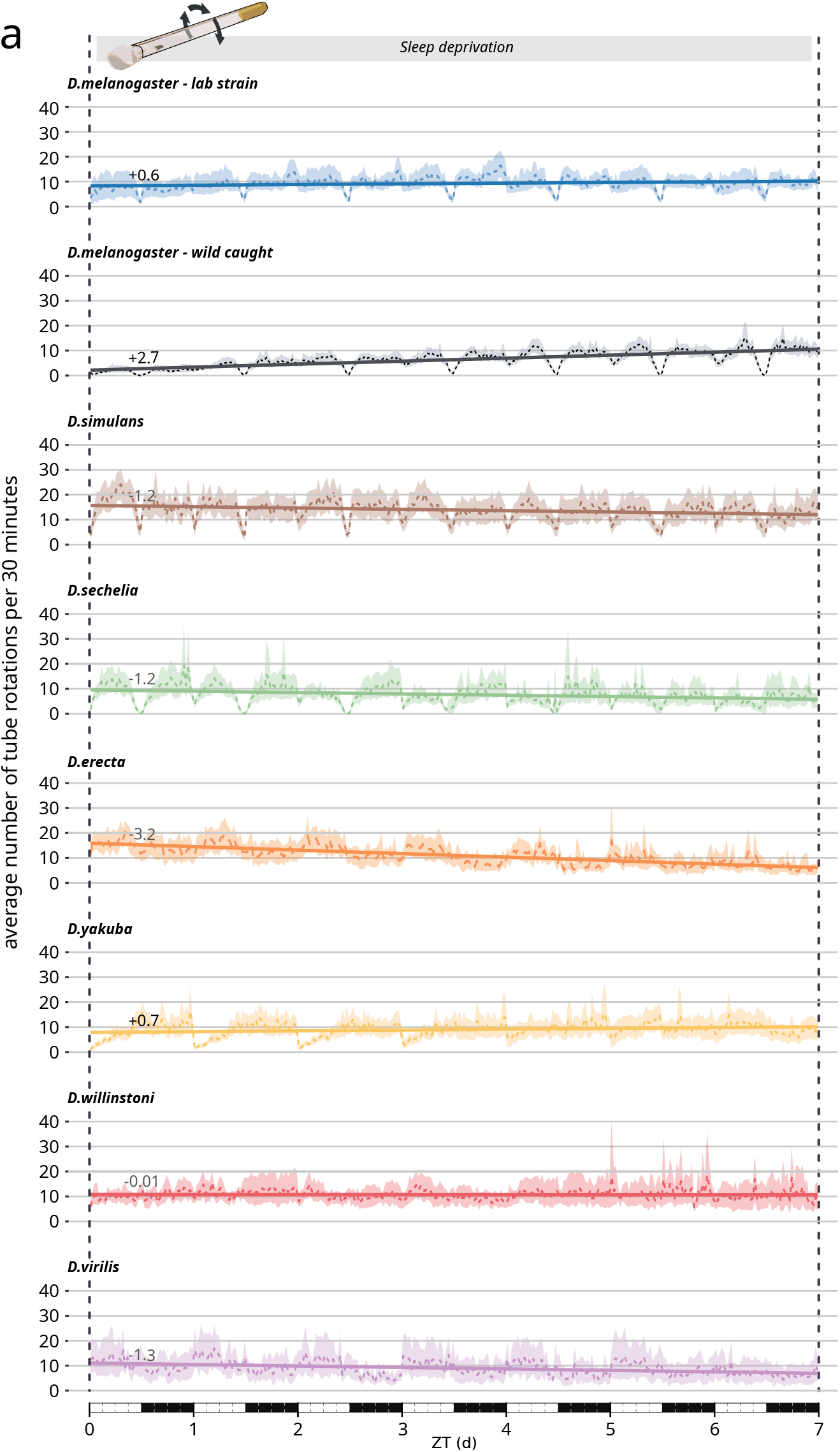
No signs of sleep pressure build-up during seven days long sleep deprivation in most species. **a**, The cumulative average number of tube rotations in 30 minutes bins in all tested species (dashed lines) and the relative trend (continuous line) calculated as time series regression linear model. The numbers above each trend line indicate the coefficient. A positive coefficient indicates a growth in sleep pressure.

Also notably, none of the species analysed showed any sign of lethality after one week of sleep deprivation (Fig. 2 supplement 4), in full accordance with earlier findings in *D. melanogaster* indicating that prolonged lack of sleep has no automatic consequences on longevity^30–32^.

**Figure 2 supplement 4.**
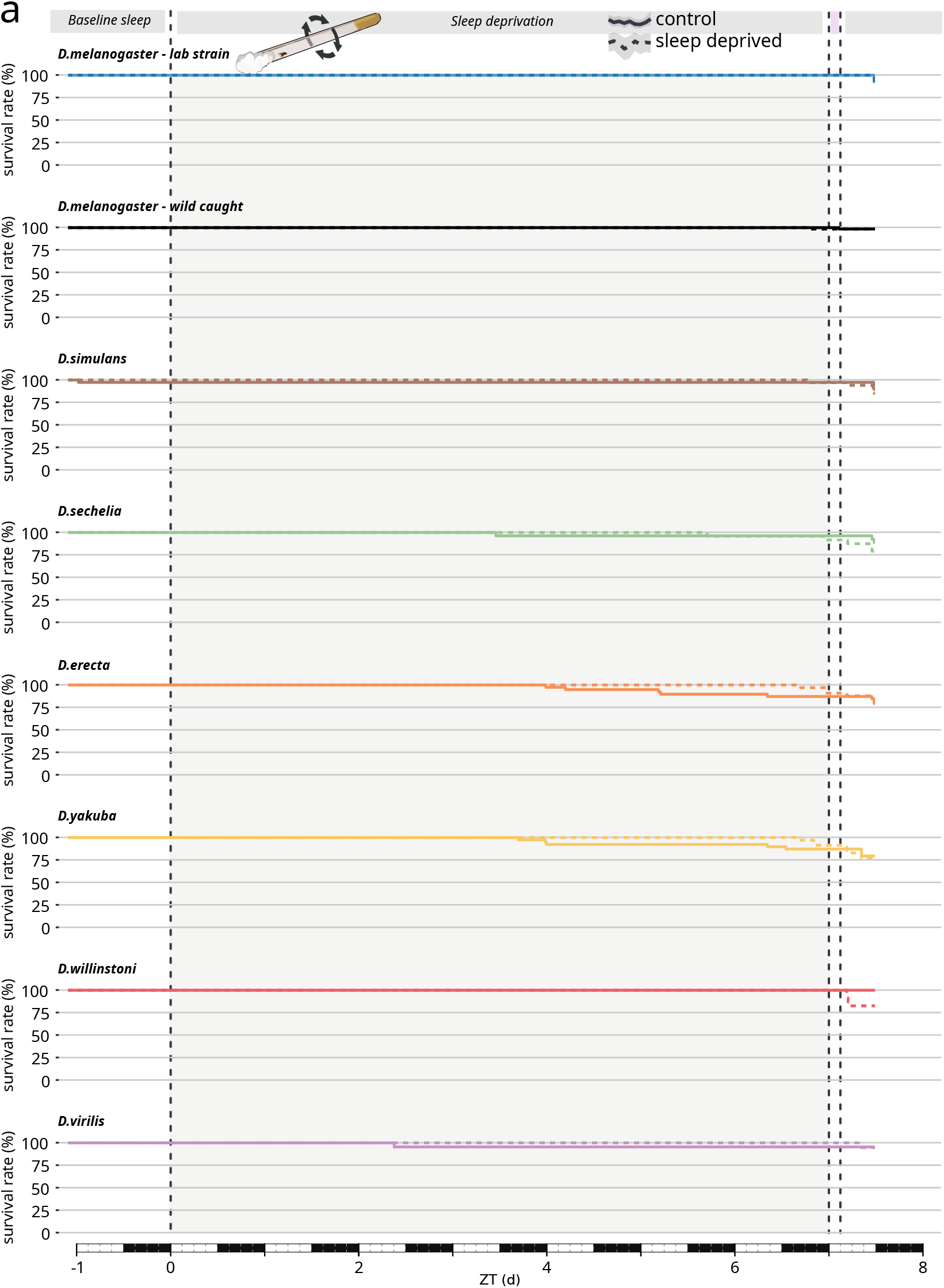
No lethality after seven days long sleep deprivation in any of the tested species. **a,** Survival curves for flies subjected to chronic, seven days long sleep deprivation, by species.

Finally, no sleep rebound could be observed even when flies were raised and maintained in condition of constant darkness (Fig. 2 supplement 5), indicating that lack of sleep rebound after sleep deprivation is not a consequence of circadian masking, but a true reflection of impaired homeostatic control.

**Figure 2 supplement 5.**
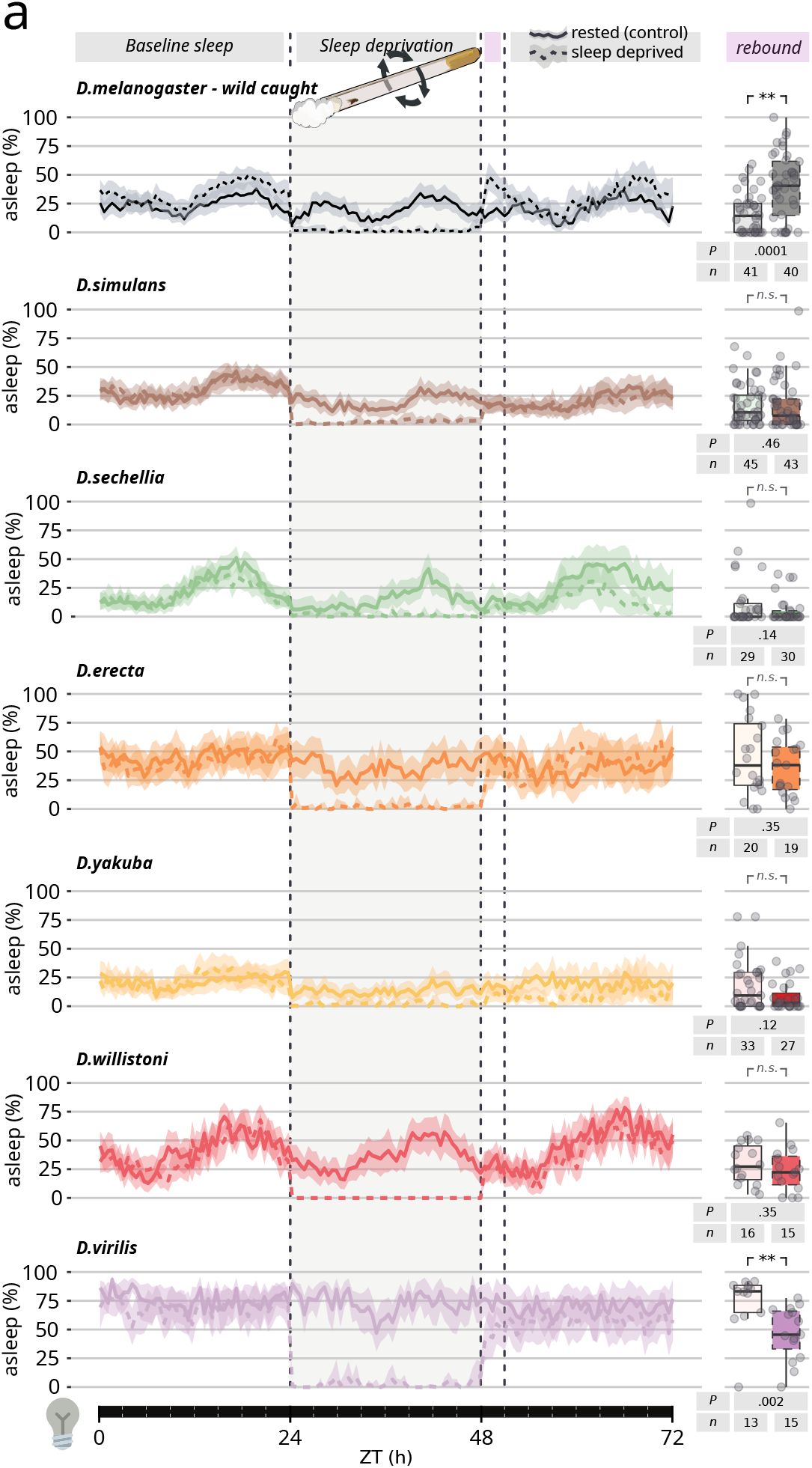
No sleep rebound, even in absence of external *zeitgebers*. **a,** 72 h sleep profile (left) and quantification of rebound (right) in flies subjected to 24 h of mechanically-induced sleep deprivation in constant dark conditions. Each panel features one strain from each of the seven wild-caught species, or CantonS. In all panels, the sleep profile of rested flies is shown as a continuous line, while sleep-deprived animals are shown in a dashed line. The sleep rebound at ZT 0-3 is quantified on the right side of each sleep profile. Numbers of animals (Ns) and P-values of sleep-deprived *vs* control are shown below each panel. Flies were raised in constant darkness too. *** P<0.001; ** P<0.01; * P<0.05.

While mechanical sleep deprivation is extremely efficient at keeping flies awake (Fig. 2 supplement 1d), its ecological relevance is debatable. We therefore explored a more natural way to keep animals awake: male-male interaction in socially naive animals^33,34^. To our surprise, a 24 hours sleep deprivation through interaction with a conspecific white-eyed male eventually led to a recognizable rebound in four of the seven tested species (Fig. 2b), all within the closest *melanogaster* subgenus. Still no rebound could be observed in *D. erecta*, *D. willistoni*, *D. virilis* (Fig. 1a).

The results of these behavioural experiments reveal different evolutionary paths of the two key aspects of sleep regulation in Drosophilids: tightly conserved circadian control, but divergent homeostatic regulation. Understanding the molecular underpinnings of these differences can help shed light not just on sleep evolution, but on the mystery of sleep homeostasis too. Work conducted in the past decades, has consolidated an emerging picture linking synaptic plasticity, learning, and experience to sleep, correlating synaptic strength to sleep needs^35–40^. We^36^ and others^37,39,41–44^ previously showed that, in *D. melanogaster*, prolonged wake is associated with a reinforcement in synaptic strength conveniently recognizable through an increase in the expression of the synaptic scaffolding protein Bruchpilot (BRP). To check whether this mechanism is common to other *Drosophila* species, we quantified the expression of BRP protein in the brains of the flies after a full day of prolonged wakefulness after mechanical (Fig. 2d) or social sleep deprivation (Fig. 2e). At least for social sleep deprivation, we found a one-to-one correlation between appearance of rebound sleep and increase in detectable BRP levels (Fig. 2e). For mechanical sleep deprivation, the correlation appeared slightly more spurious with *D. simulans* and *D. sechelia* showing an increase in BRP expression without a concomitant manifestation of sleep rebound (Fig. 2d). The other four species, however, showed no increase in rebound and no increase in BRP expression, thus confirming and reinforcing the generality of the previously postulated link between synaptic strength and sleep homeostasis^36,41^.

If the different homeostatic regulation observed in non-melanogaster species is really to be attributed to a different modulation of synaptic strength upon waking experience, then – we reasoned – it may be possible to alter *D. melanogaster* sleep regulation by interfering with its synaptic regulatory machinery, thus making *D. melanogaster*’s homeostasis resemble the homeostasis observed in other Drosophilids. To this end, we performed a targeted genetic screen using RNAi^45^ to pan-neuronally knock-down a selected panel of seven genes known to be involved in synaptic plasticity (Fig. 3a).

**Fig. 3.**
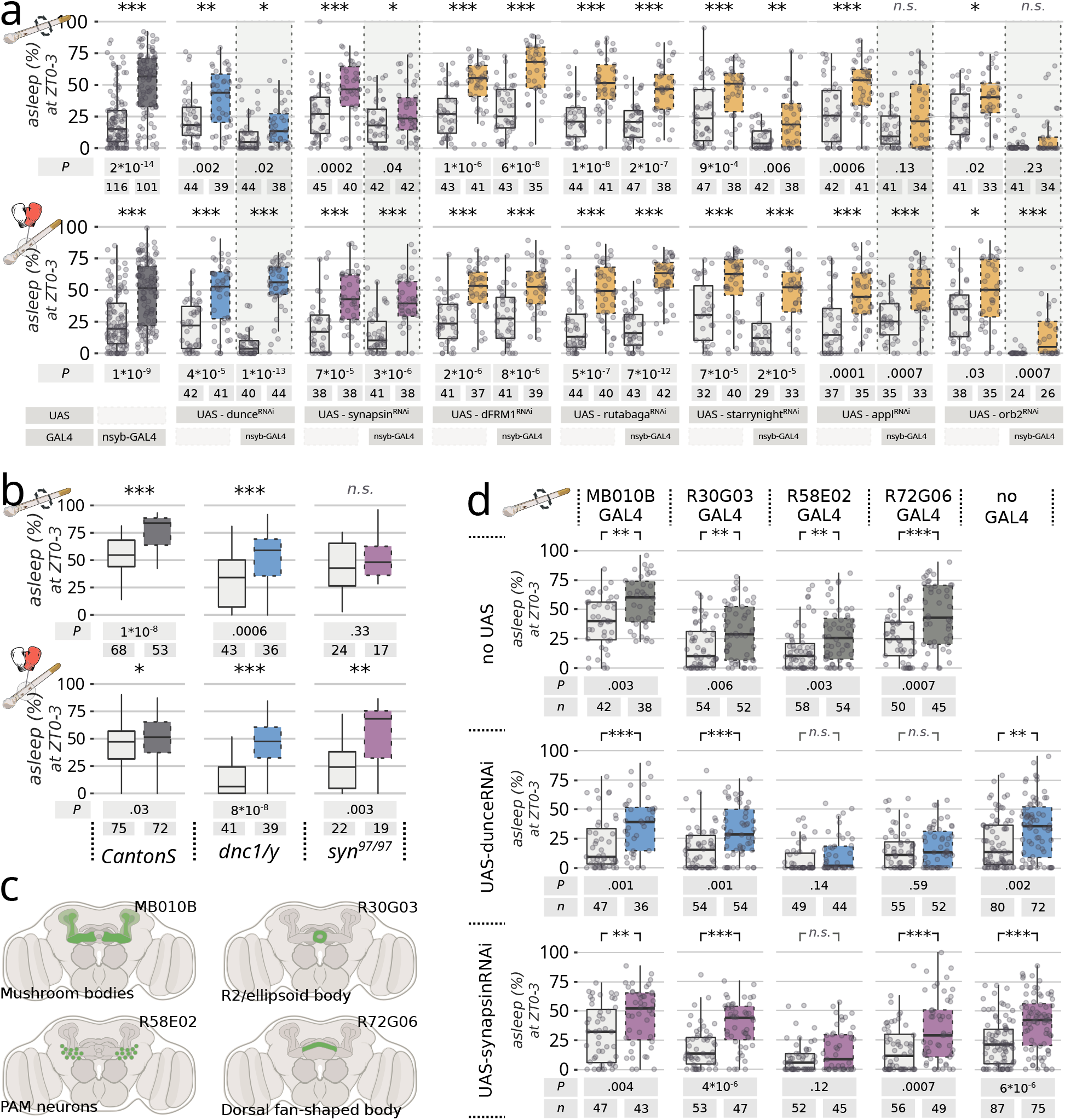
Changes in wake-induced synaptic strength can explain differences among species. **a,** Rebound sleep at ZT0-3 for flies that underwent RNAi knock-down in one of seven selected genes using the pan-neuronal GAL4 driver nSyb and their relevant parental controls. Flies underwent either 24 h of mechanical sleep deprivation (top) or 24 h of male-male sleep deprivation (bottom). **b,** Rebound sleep at ZT0-3 after 24 h of mechanical (top) or male-male (bottom) sleep deprivation in *dunce* hemizygous mutants (blue) or *synapsin* homozygous mutants (purple) compared to wild-type CantonS flies. **c,** Schematic representation of the expression of each GAL4 driver used in d. **d,** Rebound sleep at ZT0-3 for flies that underwent RNAi knock-down for *dunce* (blue) or *synapsin* (purple) in five restricted neuronal population driven by four different split-GAL4 drivers. The top-line shows the GAL4 parental control. Through the figure, sleep-deprived flies are shown with a dashed contour while rested control flies are shown in a continuous contour. Numbers of animals (Ns) and P-values of sleep-deprived *vs* control are shown below each panel. *** P<0.001; ** P<0.01; * P<0.05.

We found that, out of these seven genes, four led to a phenocopy of the non-melanogaster species when knocked-down in the brain, that is: a partial or total reduction of homeostatic rebound after mechanical sleep deprivation, but a significant rebound after male-male sleep deprivation (Fig. 3a). These were: the cAMP phosphodiesterase *dunce*^46^, the vesicle regulator *synapsin*^47^, and two genes encoding for putative amyloid-forming proteins: the *Drosophila* analogue of the β-amyloid protein precursor *appl*^48^, and the translational regulator *orb2*^49^. RNAi knock-down of those four genes either reduced (*dunce*, *synapsin*) or removed (*appl*, *orb2*) rebound after mechanical sleep deprivation without decreasing homeostatic rebound after male-male interaction, showing a behavioural phenotype analogous to the one described in *D. simulans, D. sechellia*, or *D. yakuba.* A similar effect was also observed in constitutive *synapsin* knock-out mutants (Fig. 3b). The RNAi knock-down phenotype conveniently opened the perspective of finding the neurons responsible for such a regulation, and we therefore performed a second small targeted RNAi screen looking for known sleep-regulating populations that would interfere with sleep homeostasis after sleep deprivation (Fig. 3c, d). Knock-down of *dunce* or *synapsin* in the dopaminergic PAM neurons (driven by R58E02-GAL4) and knock down of *dunce* in the dorsal fan-shaped body neurons (driven by R72G06-GAL4) interfered with the flies’ natural ability to rebound after mechanical sleep deprivation (Fig. 3d). Both regions are well known for their role in learning and memory and have been implicated with many aspects of sleep regulation^26,50–52^. These behavioural and cell-biological differences between *D. melanogaster* and other six species of the same genus suggest that spontaneous (*i.e.* circadian) sleep and homeostatic sleep evolved independently under different pressures. If this is the case, one can hypothesize that not just cellular underpinnings, but also neurochemical regulators may have followed divergent evolution.

To test this last prediction, we fed flies of all the seven tested species with chemicals acting on the two best studied neurotransmitter pathways involved in sleep regulation: chlordimeform (an octopamine^53^ agonist, Fig. 4a) and L-dopa (a dopamine^54^ precursor, Fig. 4b). Feeding flies with chlordimeform had a wake-promoting effect in all the tested species except *D. yakuba*, with the strongest arousing effect being found in *D. melanogaster*, *D. sechellia* and *D. virilis*. L-dopa, on the other hand, showed a strong wake-promoting effect in *D. melanogaster* only, with even an opposite somniferous effect in *D. sechellia* and *D. willistoni.* In Drosophilids, the neuro-regulatory dynamics of sleep have also taken divergent evolutionary paths.

**Fig. 4.**
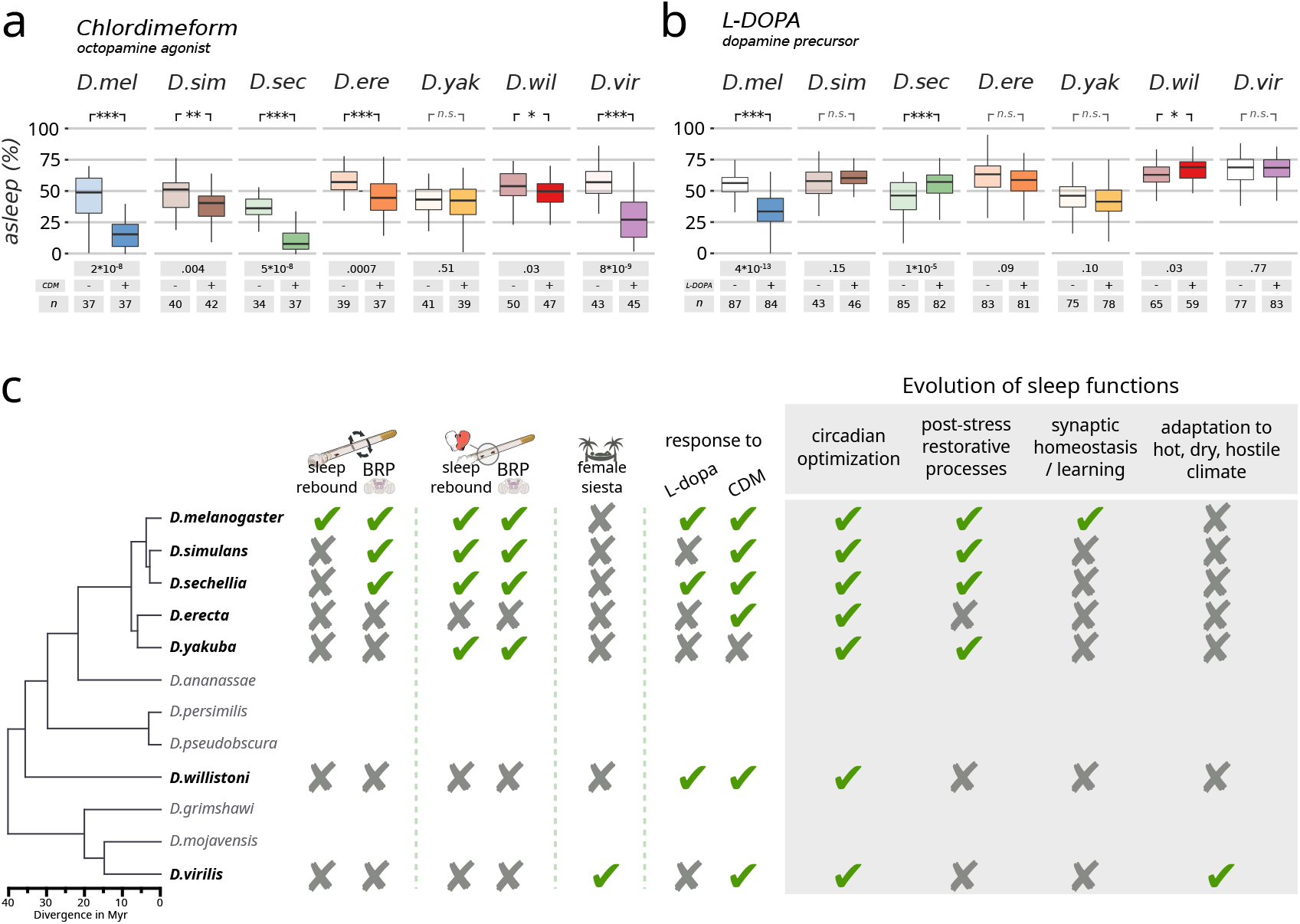
Evolutionary divergence of the neuro-pharmacology of sleep in Drosophilids. Total sleep over the 24 h period in flies fed with either chlordimeform 0.05 mg/ml, **a,** or L-DOPA 5 mg/ml, **b**. Full colours (right boxes) indicate flies fed with the drugs, while light colours (left boxes) indicate flies fed with vehicle only. Numbers of animals (Ns) and P-values are shown below each panel. *** P<0.001; ** P<0.01; * P<0.05. **c,** panoptic summary of the evolutionary divergences found in this work and the proposed model for evolution of different sleep functions (grey box).

## Discussion

Understanding the evolution of sleep in closely related species can provide unique insights into its function and regulation. One of the most elegant and powerful models used so far for this goal, is the fish *Astyanax mexicanus* which presents itself in two large conspecific populations: a long-sleeping, eyed population living in the surface of the waters, and several short-sleeping blind populations that were trapped and evolved in the darkness of Mexican caves about 2-5 million year ago, branching in an environment that provided little or no circadian *zeitgeber* and limited nutrients^5,55^. Here we introduced another powerful framework to study sleep evolution, featuring seven species of *Drosophila* spanning evolutionary distances ranging between 3 and 50 million years. To our surprise, we found that many homeostatic aspects of sleep regulation are not universally shared among these species and are, in fact, a prerogative of *D. melanogaster* only (Fig. 4c). Some (*D. simulans*, *D. sechellia*, *D. yakuba*) showed signs of sleep homeostasis only following social sleep deprivation and not after mechanical sleep deprivation, while others (*D. erecta*, *D. willistoni*, *D. virilis)* did not show any sign of homeostasis in any condition (Fig. 2a-c and Fig. 2 supplements). Sleep homeostasis is considered a defining hallmark of sleep and if we were to follow the textbook definitions^6^, we could reasonably conclude that animals showing no signs of rebound are animals that do not actually sleep: they rest, circadianly modulating their inactivity to avoid danger or maximize the return on investment of their energy expenditure. By this logic, we could argue that *D. erecta*, *D. willistoni*, *D. virilis* are non-sleeping species, but this conclusion would be at odds with the fact that all three manifested different stages of sleep depth (Fig. 1h and Fig. 1 supplement 1d). Ecologically speaking, it is easy to understand why long periods of mere inactivity could be beneficial to a well-fed animal, especially considering they happen at the riskiest times of the day (hot afternoons and dark nights)^3,4^. However, it is more difficult to explain why this relatively trivial circadian inactivity would manifest itself with different degrees of arousability. A possible explanation of this conundrum is that the phenomenon of deep sleep – whatever its function is – has evolved before its own homeostatic regulation. Based on our results, this appears to be the case in Drosophilids, and it may be a representation of a more general rule. Stages of different sleep depth in *D. melanogaster* have now been independently described by many groups^25,56–58^ and here we show they are very well conserved. Notably, we found that even their timing is conserved given that in all tested *Drosophila* species, the deepest sleep is consistently observed in the earlier part of the night – a feature well documented in mammals too.

In all species analysed in this work, the presence of sleep rebound was always accompanied by an increase in the Bruchpilot protein, an established *bona-fide* marker for synaptic strength (Fig. 2d,e). Moreover, the absence of sleep rebound could be phenocopied in *D. melanogaster* by reducing the expression of genes involved in synaptic plasticity in specific anatomical regions involved with learning or sleep (Fig. 3d). The genetic arsenal of non-melanogaster species is still budding and these latter manipulations must be limited to *D. melanogaster* for now. Nevertheless, they suggest that different dynamics of synaptic strength may explain the unexpected differences in sleep homeostasis observed among species, reinforcing the link between sleep homeostasis and synaptic strength. Importantly, genetic manipulation of synaptic plasticity specifically impaired rebound following mechanical sleep deprivation but never affected rebound following social male-male sleep deprivation (Fig. 3a,b), suggesting that the two apparently similar sleep rebounds are in fact different processes, controlled by a different machinery and possibly serving different biological functions. This would conveniently explain why some species (*D. simulans*, *D. sechelia*, *D. yakuba*) show cellular and behavioural signs of sleep homeostasis only after social male-male sleep deprivation but not after mechanical sleep deprivation. A more specific interpretation lies in the nature of the treatment itself: forced male-male interaction is likely to mimic an ecologically relevant condition of stress, which in turn has tight connections to sleep itself^59,60^. A similar paradigm, resulting in an overlapping outcome, is also been extensively studied in rodents, under the name of social defeat stress^60,61^, and it was recently shown to rely on a separate, specific neuronal circuit indeed^62^. It is tempting to speculate that the rebound sleep observed in *D. simulans*, *D. sechelia*, *D. yakuba* is driven by a specific post-stress restorative circuit, exercising a post-stree specific function and that this function might have evolved independently of other sleep functions. In other words, synaptic-homeostasis-driven sleep is different from stress-induced sleep and evolved independently of it.

Taken together, the difference in behaviour, cell-biology, and neuro-pharmacology described here, imply that the evolutionary driving force for sleep in Drosophilids is not homeostasis, as often hypothesized, but circadian adaptation. We propose that sleep in flies evolved primarily as a phenomenon to limit activity during the more dangerous or inappropriate hours of the day, restraining flies when it is too dark or too hot. All the other non-trivial functions of sleep – such as regulation of synaptic strength, learning and memory, recovery from stress, modulation of immune response, *etc.* – have then branched divergently, piggybacking on the circadian drive for inactivity (Fig. 4c) in a species-specific manner that caters diverse functions, bespoke to each species’ need. This may be the common process of sleep evolution in the animal kingdom.

## Acknowledgements

We thank the fly community for sharing reagents and protocols. Special thanks to Darren Obbard for sharing the wild-caught *D. melanogaster* strains. MJ was supported by EPSRC through doctoral grant EP/R513052/1. FF was supported by the Imperial President’s scholarship. LB was supported by BBSRC through doctoral grant BB/M011178/1. AF was supported by BBSRC through grant BB/R018839/1. L.L.P.-G.’s laboratory was supported by a European Research Council (ERC) Starting Investigator Grant (802531), a Human Frontiers Science Grant (GY0052/ 2022) and an Allen Institute Distinguished Investigator Award, as well as by The Francis Crick Institute, which is supported by core funding from Cancer Research UK (FC001594), the UK Medical Research Council (FC001594), and the Wellcome Trust (FC001594). We thank the Imperial College London Advanced Hackspace (ICAH) and the Facility for Imaging by Light Microscopy (FILM) at Imperial College London, part-supported by funding from the Wellcome Trust (grant 104931/Z/14/Z) and BBSRC (grant BB/L015129/1).

## Author-contributions

MJ, FAF, LB, ASF performed all the experiments and analysed the data with GFG; MJ, ASF, LPG and GFG devised all the experiments. GFG wrote the manuscript and all authors contributed to its editing.

## Data availability

We make publicly available all the scripts used for the data analysis (as Jupyter notebooks in Python^27^ or R^63^), all the metadata describing the experiments, and all the raw behavioural and confocal data. While the manuscript is under review, these will be available at https://lab.gilest.ro/papers/divergent-evolution-of-sleep-functions/. Upon publication everything will be frozen and made available through the Zenodo repository.

## Right assertion

For the purpose of open access, the authors have applied a Creative Commons Attribution (CC BY) licence to any Accepted Manuscript version arising, in agreement with UKRI funding requirements.

## Detailed author contributions to experimental collection and data analysis

|  |  |  |
| --- | --- | --- |
| Fig. 1 | *c* | MJ, ASF |
|  | *d,e* | MJ |
|  | *f-h* | MJ, LB |
| Fig. 1 Supplement 1 | *a-c* | MJ |
|  | *d,e* | MJ, LB |
| Fig. 2 | *a,b* | MJ |
|  | *c* | MJ, LB |
|  | *d* | MJ, FAF |
| Fig. 2 Supplement 1 | *a* | ASF |
|  | *b-d* | MJ |
| Fig. 2 Supplement 2,3,4 | *a* | MJ, FAF |
| Fig. 2 Supplement 5 | *a* | MJ |
| Fig. 3 | *a-d* | MJ |
| Fig. 4 | *a-b* | MJ, ASF |

## Materials and Methods

### Fly strains

The following VDRC-RNAi transgenic strains were used in this study: UAS-dunce^RNAi^ (#107967), UAS-synapsin^RNAi^ (#109587), UAS-dFRM1^RNAi^ (#110800), UAS-rutabaga^RNAi^ (#101759), UAS-starrynight^RNAi^ (#107993), UAS-appl^RNAi^ (#108312) and UAS-orb2^RNAi^ (#11753). The nSyb-GAL4 and MB010B-GAL4 were gifted to us by Crystal Vincent (Imperial College London,UK) and Andrew Lin (The University of Sheffield,UK) respectively, while the other GAL4 lines were obtained from Bloomington Drosophila Stock Centre (BDSC, Indiana, USA): R30G03-GAL4 (#49646), R52B10-GAL4 (#69657), R58E02-GAL4 (#41347), R72G06-GAL4 (#39792). All RNAi and GAL4 lines were outcrossed for 6 generations to the W^1118^ background before testing. The dunce^1^ (#6020), synapsin^97^ (#29031), OregonR (#5) and W^1118^ (#3605) flies, were also obtained from BDSC; CantonS strain originally from Ralf Stanewsky (Münster University, Germany). The two *D. melanogaster* wild-caught strains were a gift of Darren Obbard (University of Edinburgh, UK). Two strains of non-melanogaster species were used in Fig. 1C, and the first strain of each were studied in the succeeding figures: *D.simulans* (14021-0251.254 #60, 14021-0251.196 #61), *D.sechellia* (14021-0248.25 #3, 14021-0248.28 #53), *D.erecta* (14021-0224.01 #11), *D.yakuba* (14021-0261.01 #5, 14021-0261.48 #51), *D.willistoni* (14030-0811.24 #1, 14030-0811.13 #55) and *D.virilis (*15010-1051.87 #9, 15010-1051.118 #54). The wildtype species and the white-eyed ‘intruder’ species: *D.simulans w^-^* (14021-0251.133), *D.sechellia w^-^ (*14021-02048.30*), D.erecta w^-^ (*14021-0248.30), *D.yakuba w- (*14021-0261.04), *D.willistoni w-* (14030-10811-33) and *D.virilis w- (*15010-1051.45) were acquired from The National *Drosophila* Species Stock Center (NDSSC, Cornell University, USA).

### Fly rearing

Flies were raised on standard polenta and yeast based fly media (agar 96g, polenta 240g, fructose 960g, Brewer’s Yeast 1200g in 12 litres of water), referred to as the standard food diet or supplemented with a layer of potato starch (66-117, Nutri-Fly® Instant) (1:4 potato starch:H_2_O) and filter paper (referred as: the potato starch diet). The species panel were reared on the potato starch diet in all but Fig. 2, where the flies were raised in standard food. The knockdown and mutant data in Fig. 3B-D were raised on the standard food diet. The species food was supplemented with potato starch to improve their viability and filter paper to encourage egg laying and increase progeny number.

### Behavioural experiments

#### Sleep analysis in ethoscopes

For all experiments, 0-3 days male or virgin female adult flies were sorted into glass tubes (70×5×3 mm [length × external diameter × internal diameter]) containing standard food. The tubes were loaded into ethoscope sleep arenas (20 animals per device). At least one day of baseline was recorded before any treatment. All experiments were carried out under LD conditions unless stated otherwise, 50-70% humidity, in incubators set at 25°C and with *ad libitum* access to regular food. Animals that died during the experiment were excluded from the analysis. In order to avoid confounding effects related to the location of the tube on sleep amount (*e.g.* an ethoscope and incubator), the position of all the tubes was systematically interspersed. The endogenous period length was established with chi-square χ^2^ periodic analysis*^1^*averaged over 5 days of activity in constant conditions. Flies were both raised and tested with a DD light cycle.

#### Mechanical Sleep deprivation

The effects of mechanical sleep deprivation shown in Fig. 2a were tested using the optomotor module*^2^*, programmed with a 30 seconds immobility trigger. When triggered, a motor rapidly turns (∼400 rpm) for 3 seconds, spinning the tube housing the fly and preventing sleep. In Fig.3 the motor stimulus had a duration of 1 second. The mechanical sleep deprivation typically lasted 24 hours, except in the case of the continuous mechanical sleep deprivation where it was programmed to sleep deprive the fly for 168 hours. Response to a reduced stimulus of ∼150 rpm (Fig. 2 supplement 1a) were tested with the servo module*^3^*. Animals recovered for 1 day in the ethoscopes to measure rebound. Animals that were asleep >30% of the sleep deprivation period were excluded from the analysis.

For the mechanical rebound response in DD (Fig. 2 supplement 5), flies were raised and tested in constant darkness with 6 days of baseline, 24hr mechanical sleep deprivation and 1 day recovery for rebound measurement.

#### Male-male sleep deprivation

For social interactions, flies were removed from a shared vial and placed in 70 mm x 5 mm glass tubes containing standard food. Twenty tubes were placed in each ethoscope arena. Flies were acclimated in behavioural glass tubes for 4 days of which the last 1-2 days were recorded as a baseline. On the interaction day, intruders (conspecific white-eyed males) were added at ZT0. Intruders were then removed in darkness (ZT23-ZT24). Rebound period was then recorded. All figures show the last baseline day and the first rebound day.

#### Analysis of sleep depth

Arousal threshold during sleep baseline was assessed empirically using the air stimulation module of the ethoscope platform (motorized air/gas/odour delivery module or mAGO)*^3^*. After two days of baseline sleep recording, ethoscopes were programmed to deliver air stimuli with a 30s immobility trigger, using a stimulus delivery probability factor of 10% to allow for mock stimulation to be used as control, and to extend the bout length of disturbed inactivity to several multiples of 30s. A second similar experiment was conducted with an 150s immobility trigger and 50% probability factor. After each experiment, behavioural data were analysed using species-specific hidden Markov chain models in ethoscopy, as described*^4^*. For analysis of sleep states during rebound, the species specific hidden Markov chain models were applied to the unprocessed position data from the ethoscopes to investigate the time spent in deep sleep, light sleep, quietly awake and fully awake after 24hr mechanical sleep deprivation.

#### Behavioural response to pharmacological stimulation

For the drug treatments, L-dopa (5 mg/ml) was added to sucrose-based food (1% Agar, 5% Sucrose) while the CDM (0.05 mg/ml) was prepared in standard food. One day of baseline behaviour was recorded on a control diet (complementary diet minus drug) before flies were flipped onto the drug treatment for 48 hours. Successful food consumption was assessed using a blue coloured dye*^5^*.

### Immunohistochemistry

Brains were dissected and stained as described*^6^*. The species panel were aged to 10 days in a 25°C incubator before setting up for 24 hours mechanical or social sleep deprivation. At ZT0, on the morning after sleep deprivation, flies were immobilized on ice for brain dissection in 0.1M phosphate buffered saline (PBS). The brains were fixed in 4% paraformaldehyde for 20 minutes at room temperature before performing 3 x 10 minutes washes in PBST (0.3% Triton X in PBS). Samples were then blocked at room temperature for 1 hour in 5% normal goat serum (NGS) in PBST followed by incubation with 1:10 mouse anti-nc82 (AB_2314866, DSHB) for 48 hours at 4°C. After primary antibody incubation, the brains were washed for 3 x 10 minutes in PBST and incubated with 1:200 goat anti-mouse IgG (ab150115, abcam) in 5 % NGS for 48 hours at 4°C. Another 3 x 10 minute washes in PBST were completed before mounting brains on a microscope slide with vectashield (Vector Laboratories) for imaging. Brains were imaged with a Leica SP8 - STELLARIS 5 Inverted Light Sheet Confocal Microscope at x40 magnification to capture z stack images of the dorsal and ventral brain regions. For comparative analysis of the expression levels, images were taken in the same confocal session using identical laser and confocal settings. Analysis of the data was performed using Fiji/Image J*^7^* (NIH, Bethesda). The fan-shaped body (FSB) region was manually delimited stack by stack and a maximum intensity projection was generated. Background measurements were taken from the image background around the brain and subtracted to generate the intensity values. The data was normalised within repeats.

### Statistics, Data availability, and Reproducibility

All ethoscope data were analysed using rethomics*^8^* or ethoscopy*^4^*. Statistical comparisons were performed as indicated in the text and figure legends, mostly using Wilcoxon rank sum test with false rate discovery (FDR) correction. In all summary plots, the intermediate reference mark indicates the median and the surrounding error estimates always indicate the bootstrapped 95% confidence intervals. Whenever possible, the entire dataset is shown as a dot plot. All figures explicitly mention the biological Ns *i.e.* the number of biologically independent animals for each data point. Each conclusion relies on multiple independent experiments and never fewer than three independent ones. The actual number of experiments for each panel can be found in the metadata descriptions that are supplied along with the R and Python scripts. Unless differently stated in the legend, all P-values arise from Wilcoxon rank sum tests. P-values are intended to be supportive and indicate where statistical significance occurs in presence of slight CI limit overlap. In all figures, * are used to indicate customary thresholds of statistical significance: P<0.05 *; P<0.01 **; P<0.001 ***. The actual numerical P-value is shown in each figure whenever possible and full statistical comparisons among all combinations are available as extended data information in a dedicated file. Moreover, all the scripts (in R and Python) used to generate the figures in this manuscript as well the related statistical analysis and the original behavioural raw data as obtained with ethoscopes are publicly available on the Gilestro laboratory website https://lab.gilest.ro/papers/divergent-evolution-of-sleep-functions/ and will be frozen and made available through the Zenodo repository upon formal pubblication. All the hardware and software created in the lab is open source and can be explored at http://lab.gilest.ro/ethoscope3 and http://lab.gilest.ro/rethomics8. Rethomics versions used to analyse the data were as follows: behavr: 0.3.2; sleepr: 0.3.0; zeitgebr: 0.3.3; ggetho: 0.3.4; scopr: 0.3.3.

